# Interaction of the spike protein RBD from SARS-CoV-2 with ACE2: similarity with SARS-CoV, hot-spot analysis and effect of the receptor polymorphism

**DOI:** 10.1101/2020.03.04.976027

**Authors:** Houcemeddine Othman, Zied Bouslama, Jean-Tristan Brandenburg, Jorge da Rocha, Yosr Hamdi, Kais Ghedira, Najet Srairi-Abid, Scott Hazelhurst

## Abstract

The spread of COVID-19 caused by the SARS-CoV-2 outbreak has been growing since its first identification in December 2019. The publishing of the first SARS-CoV-2 genome made a valuable source of data to study the details about its phylogeny, evolution, and interaction with the host. Protein-protein binding assays have confirmed that Angiotensin-converting enzyme 2 (ACE2) is more likely to be the cell receptor through which the virus invades the host cell. In the present work, we provide an insight into the interaction of the viral spike Receptor Binding Domain (RBD) from different coronavirus isolates with host ACE2 protein. By calculating the binding energy score between RBD and ACE2, we highlighted the putative jump in the affinity from a progenitor form of SARS-CoV-2 to the current virus responsible for COVID-19 outbreak. Our result was consistent with previously reported phylogenetic analysis and corroborates the opinion that the interface segment of the spike protein RBD might be acquired by SARS-CoV-2 via a complex evolutionary process rather than a progressive accumulation of mutations. We also highlighted the relevance of Q493 and P499 amino acid residues of SARS-CoV-2 RBD for binding to human ACE2 and maintaining the stability of the interface. Moreover, we show from the structural analysis that it is unlikely for the interface residues to be the result of genetic engineering. Finally, we studied the impact of eight different variants located at the interaction surface of ACE2, on the complex formation with SARS-CoV-2 RBD. We found that none of them is likely to disrupt the interaction with the viral RBD of SARS-CoV-2.

## 1 Introduction

The coronavirus SARS-CoV-2 (previously known as nCoV-19) has been associated with the recent epidemic of acute respiratory distress syndrome [2]. Recent studies have suggested that the virus binds to the ACE2 receptor on the surface of the host cell using spike proteins, and explored the binary interaction of these two partners [8, 23]. In this work, we focused our analysis on the interface residues to get insight into four main subjects: (1) The architecture of the spike protein interface and whether its evolution in many isolates supports an increase in affinity toward the ACE2 receptor; (2) How the affinity of SARS-COV-2-RBD and SARS-CoV-RBD toward different ACE2 homologous proteins from different species is dictated by a divergent interface sequences (3); A comparison of the interaction hotspots between SARS-CoV and SARS-CoV-2; and finally, (4) whether any of the studied ACE2 variants may show a different binding property compared to the reference allele. To tackle these questions we used multi-scale modelling approaches in combination with sequence and phylogenetic analysis.

## 2 Materials and Methods

### 2.1 Sequences and data retrieval

Full genome sequences of 10 Coronaviruses isolates were retrieved from NCBI Genbank corresponding to the following accession numbers: AY485277. (SARS coronavirus Sino1-11), FJ882957.1 (SARS coronavirus MA15), MG772933.1 (Bat SARS-like coronavirus isolate bat-SL-CoVZC45), MG772934.1 (Bat SARS-like coronavirus isolate bat-SL-CoVZXC21), DQ412043.1 (Bat SAR coronavirus Rm1), AY304488.1 (SARS coronavirus SZ16), AY395003.1 (SARS coronavirus ZS-C), KT444582.1 (SARS-like coronavirus WIV16) MN996532.1 (Bat coronavirus RaTG13) in addition to Wuhan seafood market pneumonia virus commonly known as SARS-CoV-2 (accession MN908947.3).

The sequences of the surface glycoprotein were extracted from the Coding Segment (CDS) translation feature from each genome annotation or by locally aligning the protein from SARS-CoV-2 with all possible ORFs from the translated genomes. ACE2 orthologous sequences from Human (Uniprot sequence Q9BYF1), Masked palm civet (NCBI protein AAX63775.1 from *Paguma larvata*), Chinese rufous horseshoe bat (NCBI protein AGZ48803.1 from *Rhinolophus sinicus*), King cobra snake (NCBI protein ETE61880.1 from *Ophiophagus hannah*), chicken (NCBI protein XP 416822.2, *Gallus gallus*), domestic dog (NCBI protein XP 005641049.1, *Canis lupus familiaris*), pig (NCBI protein XP 020935033.1, *Sus scrofa*) and Brown rat (NCBI protein NP 001012006.1 *Rattus norvegicus*) were also computed and retrieved.

Human variants of the ACE2 gene were collected from the gnomAD database. Only variants that map to the protein coding region and belonging to the interface of interaction with the RBD of the spike protein were retained for further analyses.

### 2.2 Sequence analysis and phylogenetic tree calculation

MAFFT 7.450 was used to align the whole genome sequences and the protein sequences of viral RBDs [5] (Supplementary Materials 1). Prediction of the N-Glycosylation sites was made for all studied ACE2 sequences using NetNGlyc server (https://www.cbs.dtu.dk/services/NetNGlyc/). For the genome comparison, we selected the best site model based on lowest Bayesian Information Criterion (BIC) calculated using model selection tool implemented in MEGA 6 software [16]. The General Time Reversible (GTR) model was chosen as the best fitting model for nucleotide substitution with discrete Gamma distribution (+G) with 5 rate categories. For the RBD sequences, the best substitution model for maximum likelihood (ML) calculation was selected using a model selection tool implemented on MEGA 6 software based on the lowest BIC score. Therefore, the WAG model [20] using a discrete Gamma distribution (+G) with 5 rate categories has been selected.

Phylogenetic trees were generated using a ML method in MEGA 6. The consistency of the topology, for the RBD sequences, was assessed using a bootstrap method with 1000 replicates. The resulting phylogenetic tree was edited with iTOL [9].

### 2.3 Homology based protein-protein docking and binding energy scores estimation

The co-crystal structure of the spike protein of SARS-CoV complexed to human-civet chimeric receptor ACE2 was solved at 3 *Å* of resolution (PDB code 3SCL). We used this structure as a template to build the complex of spike protein from different virus isolates with the human ACE2 protein (Uniprot sequence Q9BYF1). The template sequences of the ligand (spike protein) and the receptor (ACE2) were aligned locally with the target sequences using the program Water from the EMBOSS package [12]. Modeller version 9.22 [14] was then used to predict the complex model of each spike protein with the ACE2 using a slow refining protocol. For each model, we generated ten conformers from which we selected the model with the best DOPE score [15].

To calculate the binding energy scores we used, PRODIGY server [22], MM-GBSA method implemented in the HawkDock server [19] and FoldX5 [3]. The contribution of each amino acid in protein partners was calculated HawkDock server. Different 3D structures of human ACE2 (hACE2), each comprising one of the identified variants, were modeled using the BuildModel module of FoldX5. Because it is more adapted to predict the effect of punctual variations of amino acids, we used DynaMut at this stag of analysis [13].

### 2.4 Flexibility analysis

We ran a protocol to simulate the spike RBD fluctuation of SARS-CoV- and SARS-CoV using the standalone program CABS-flex (version 0.9.14) [7]. Three replicates of the simulation with different seeds were conducted usin a temperature value of 1.4 (dimensionless value related to the physica temperature). The protein backbone was kept fully flexible and the numbe of the Monte Carlo cycles was set to 100.

## 3 Results

### Sequence and phylogenetic analysis

Phylogenetic analysis of the different RBD sequences revealed two wel supported clades. Clade 1 includes Rm1 isolate, Bat-SL-CoVZC45 an Bat-SL-CoVZXC21. These three isolates are closely related to SARS-CoV-as revealed by the phylogenetic tree constructed from the entire genom (Figure 1A). Clade 2 includes SARS-CoV-2, RatG13, SZ16, ZS-C, WIV16 MA15, and SARS-CoV-Sino1-11 isolates (Figure 1A). SARS-CoV-2 an RatG13 sequences are the closest to the common ancestor of this clad The exact tree topology is reproduced when we used only the RBD segmen corresponding to the interface residues with hACE2. This is a linea sequence spanning from residue N481 to N501 in SARS-CoV-2.

**Figure 1.**
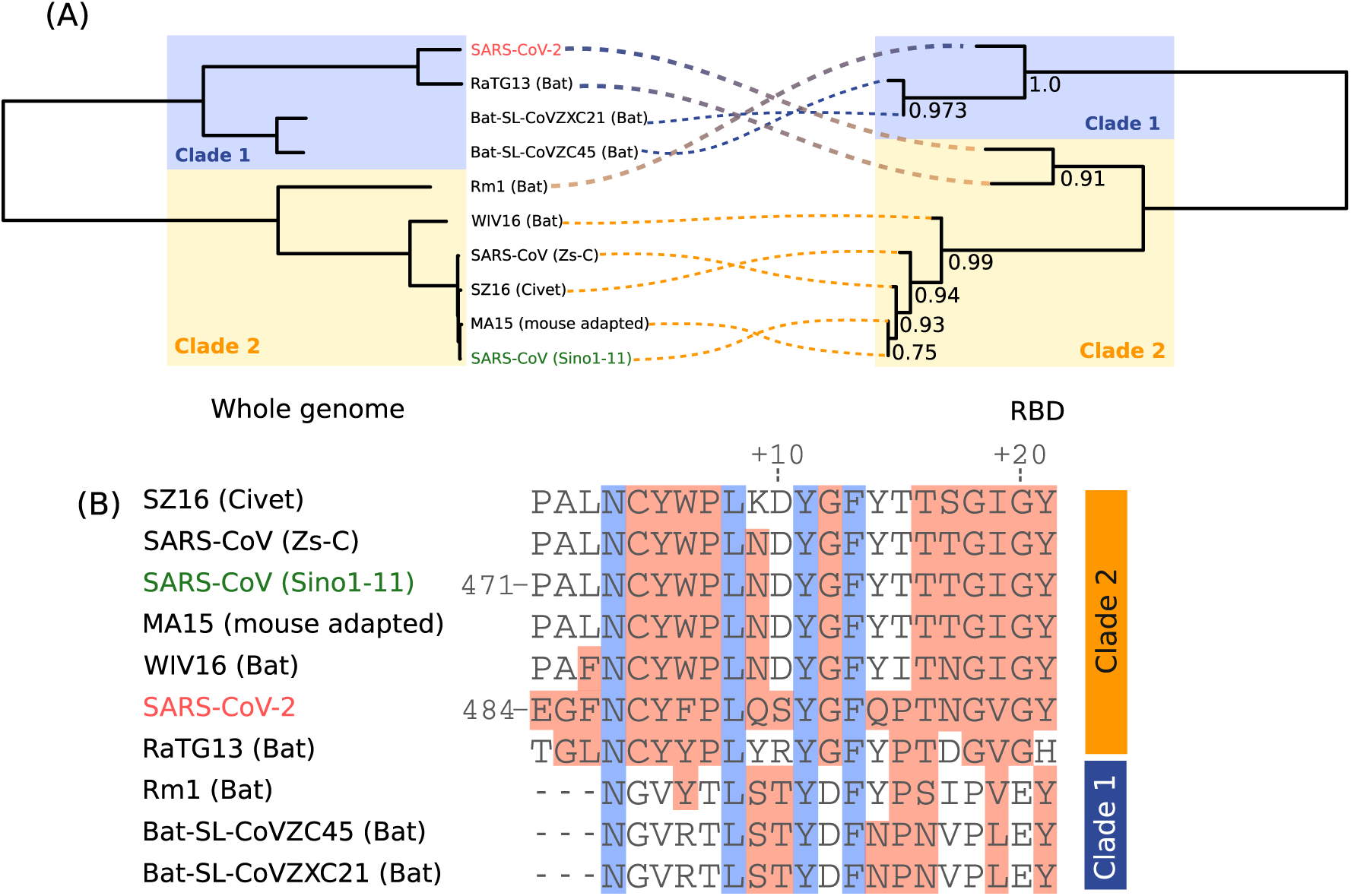
Phylogenetic and sequence analysis based on full genomes and RBDs from the different isolates included in this study. (A) Phylogeny trees are opposed to each other to show the clade discrepancies and discontinuous lines shows the equivalent taxon between each tree. (B) Multiple sequence alignment of the interface residues of RBD. Blocks in red color indicate the residues with similar biochemical properties to the positions in SARS-CoV-2. Conserved residues are colored in blue.

Multiple sequence alignment showed that the interface segment of SARS CoV-2 shares higher similarity to sequences from clade 2 (Figure 1B However, we noticed that S494, Q498 and P499 are exclusively simila to their equivalent amino acids in sequences from clade 1. SARS-CoV-interface sequence is closely related to RaTG13 sequence, isolated from *Rhinolophus affinis* bat.

### 3.1 Prediction of the RBD/hACE2 complex structure

To investigate whether the interface of the spike protein isolate evolves by increasing the affinity toward the ACE2 receptor in the final host, we predicted the interaction models of the envelope anchored spike protein (SP) from several clinically relevant coronavirus isolates with hACE2 receptor (PDB files for the complexes are listed in Supplementary Materials 1). The construction of the complex applies a comparative-based approach that uses a template structure in which both partners (ligand and receptor) are closely related to those in the target system respectively. In our study, we only modeled the interaction of the RBD which was shown to be implicated in the physical interaction with ACE2 (Figure 2A). The lowest sequence identit of the modeled spike proteins as well as those of any of the orthologou ACE2 sequences (Human, civet, bat, pig, rat, chicken and snake) do not fal below 63% toward their respective templates. At such values of sequenc identities it is expected that the template and the target complexes shar the same binding mode [6].

**Figure 2.**
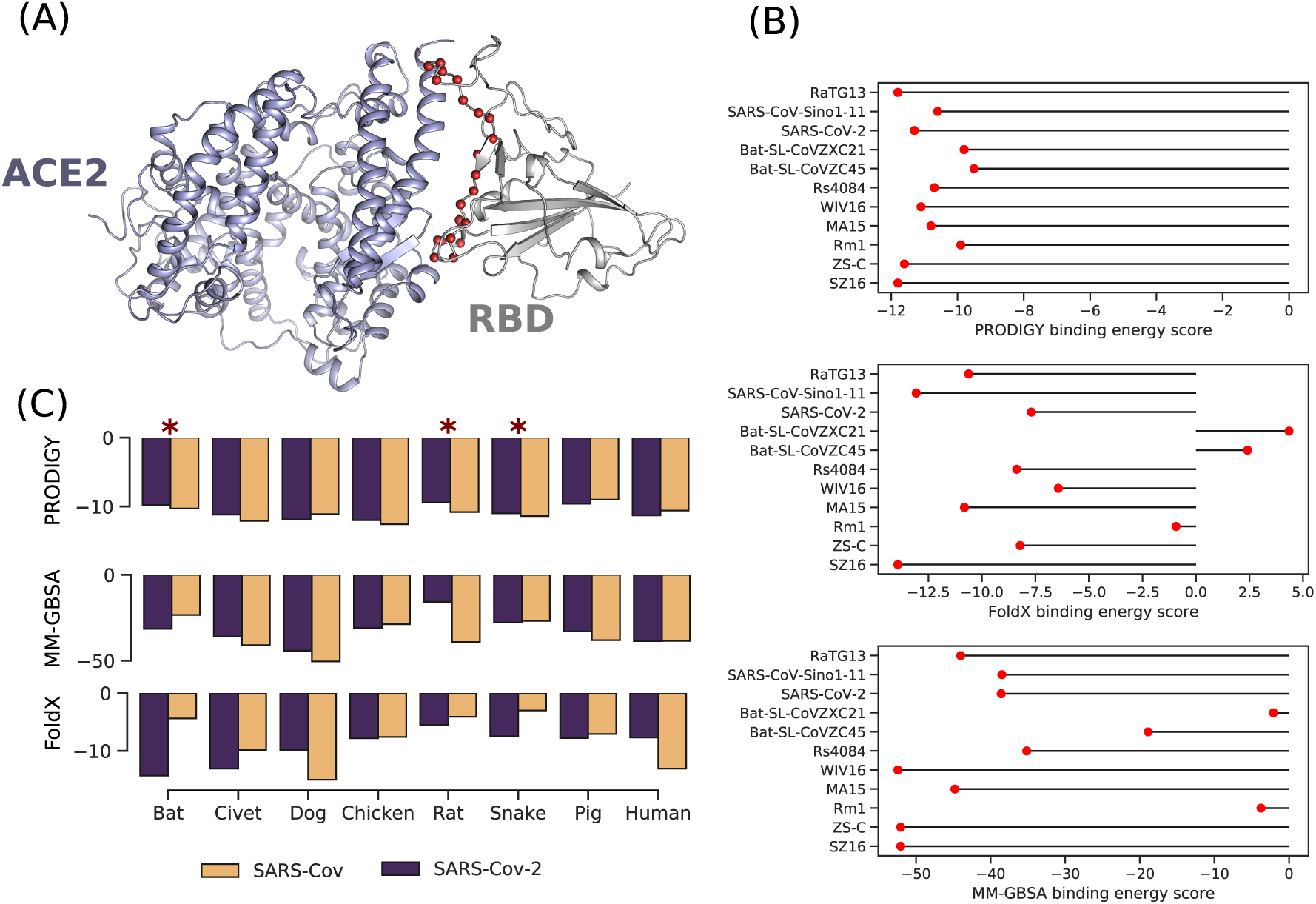
Homology based protein-protein docking of RBD/ACE2 and binding energy score analysis of spike RBD with ACE2 receptor. (A) Homology based protein-protein docking complex of SARS-CoV-2 RBD with hACE2. The red spheres are the interface residues of the RBD. (B) Binding energy scores calculated with PRODIGY, MM-GBSA and FoldX methods for RBDs from different coronaviruses forms with hACE2. (C) Binding energy scores of RBDs from SARS-CoV-2 and SARS-CoV interacting with ACE2 orthologues. Asterisks indicate the putative overlap of a glycosylation site with the protein-protein interface.

### 3.2 Analysis of the interaction energy scores of hACE with other virus isolates

We calculated the binding energy scores of the RBD from different viru isolates interacting with hACE2 (Figure 2b). All three methods use for the calculation are in agreement that RBDs from bat-SL-CoVZC45 bat-SL-CoVZXC21 and Rm1 show the worst energy scores. While the binding energy score falls in the boundary limit of the incertitude margin fo PRODIGY calculation (section 2, Supplementary material 2), the difference in the scores calculated by FoldX and MM-GBSA are not. Therefore w consider that such differences in energies compared to SARS-CoV-2 are consistent between the three methods. Except for FoldX, the affinity in predicted to be more favorable for RBD from SARS-CoV-2 compared t SARS-CoV. However, MM-GBSA only marginally discriminates betwee the two values.

### 3.3 Interaction of RBD from SARS-CoV-2 and SAR CoV with different ACE2 orthologues

To investigate the tendency of SARS-CoV-2 and SARS-CoV to interac with different orthologous forms of ACE2, we analysed the divergence in their respective interacting surfaces. We have also mapped the putativ glycosylation sites that overlap with the interface with RBD. Overall, the binding energy scores are similar between SARS-CoV-2 and SARS-Co considering the estimation of error for each method. Variances are mor important for the calculations made by FoldX and although of differen formalism, MM-GBSA and PRODIGY scores are relatively in agreemen Compared to hACE2, only the Canidae form shows better energy score both in PRODIGY and MM-GBSA for SARS-CoV-2. Moreover, We found that putative glycosylation sites overlap significantly with RBD interaction in Snake, Rat and Bat forms (section 3, Supplementary data 2).The docking also shows that key residues of RBD SARS-CoV-2 tend to interact with conserved residues on ACE2 (Figure 3, Supplementary data 2) (residues 36-53 in hACE2) which can explain the similar values of energy scores.

**Figure 3.**
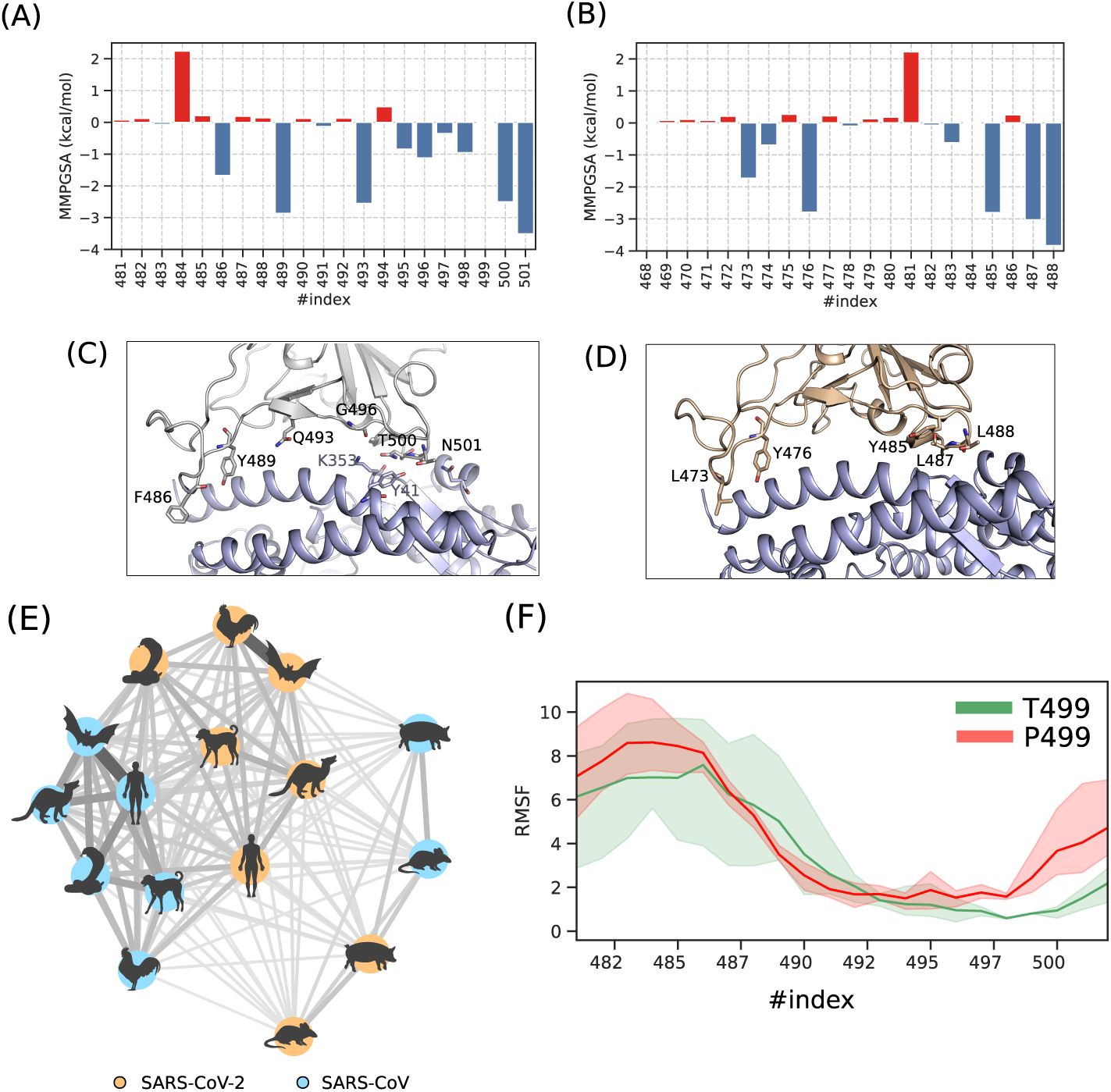
Analysis of the interaction between RBD and hACE2. Decomposition of the MM-GBSA energy for each amino acid of the binding surface from SARS-CoV-2 (A) and SARS-CoV Sino1-11 isolate (B). Position of the hotspot residues of the complexes RBD-SARS-CoV-2/hACE2 (C) and RBD-SARS-CoV/hACE2 (D). (E) similarity matrix in network representation calculated from the free energy decomposition profiles of complexes involving SARS-CoV-2 and SARS-CoV RBDs interacting with different orthologous sequences of ACE2. (F) Flexibility of RBD interface residue expressed as Root Mean Square Fluctuation (RMSF) for two forms of RBD-SARS-CoV-2, T499 and P499.

### 3.4 Decomposition of the interaction energy

MM-GBSA allowed us to assign the contribution of each amino acid in the interface with hACE2, in the binding energy score. We conducted this analysis using both sequences of the SARS-CoV-2 Wuhan-Hu-1 (Figure 3A and the Sino1-11 SARS-CoV (Figure 3B) isolates. Residues F486, Y489, Q493, G496, T500 and N501 of SARS-CoV-2 RBD forming the hotspots of the interface with hACE2 protein were investigated (we only consider values > 1 or < 1 kcal/mol to ignore the effect due to the thermal fluctuation). All these amino acids form three patches of interaction spread along the linear interface segment (Figure 3C): two from the N and C termini and one central. T500 establishes two hydrogen bonds using its side and main chains with Y41 and N330 of hACE2. N501 forms another hydrogen bond with ACE2 residue K353 buried within the interface. On the other hand, SARS-CoV RBD interface contains five residues (Figure 3D), L473, Y476, Y485, T487 and T488 corresponding to the equivalent hotspot residues of RBD from SARS-CoV-2 F487, Y490, G497, T501 and N502. Therefore, Q493 as a hotspot amino acid is specific to SARS-CoV-2 interface. The equivalent residue N480 in SARS-CoV only shows a non-significant contribution of 0.18 kcal/mol.

The similarity matrix analysis was conducted to assess the divergence of the interaction interface of RBDs qualitatively, i.e. the specific set of residues implicated in the interaction with ACE2, and quantity, i.e. the contribution of each residue in the binding energy score. The similarity matrix was calculated from free energy decomposition of interface residues of RBDs from SARS-CoV-2 and SARS-CoV in complex with ACE2 orthologous and reported as a network representation (Figure 3E and Figure 1 and 2 in Supplementary Materials 2). We noticed the existence of densely interconnected edges involving all the protein-protein complexes for SARS-CoV-2 and SARS-CoV except those involving ACE2 from *Sus scrofa* and *Rattus norvegicus*. Complexes involving the RBD of SARS-CoV-2 show less intrinsic similarity compared to RBD of SARS-CoV. However, similarity scores tend to be uniform in the group involving ACE2 from human, civet, dog, bat, snake, and chicken. The complex including hACE2 does not seem to diverge from the rest of the members of the SARS-CoV-2 group such as the case of *Sus scrofa* and *Rattus norvegicus*.

### 3.5 Flexibility analysis

Sequence analysis and the visual inspection of RBD/hACE2 complex might reflect the substitution of P499 in SARS-CoV-2 RBD as a form of adaptation toward a better affinity with the receptor. In order to further investigate its role, we performed a flexibility analysis using a reference structure (SARS-CoV-2 RBD containing P499) and an *in silico* mutated form P499T, a residue found in SARS-CoV and most of the clade 2. Our results show that the mutation caused a significant decrease in stability for nine residues of the interface corresponding to segment 482-491 (Figure 3F). Indeed, the RMSF variability per amino acid for this sequence increases compared to the reference structure.

### 3.6 Analysis of ACE2 variability and affinity with the virus

A total of eight variants of hACE2 that map to the interaction surface are described in the gnomAD database (Figure 4A). All these variants are rare (Table 1) and mostly found in European non-Finnish and African populations. Considering both the enthalpy (*ddG*) and the vibrational entropy in our calculation (*ddS*), we found no significant changes (> 1 or < 1 kcal/mol) in neither the folding energy of the complex (Figure 4B) nor the interaction energy of the protein-protein partners (Figure 4C).

**Table 1.**
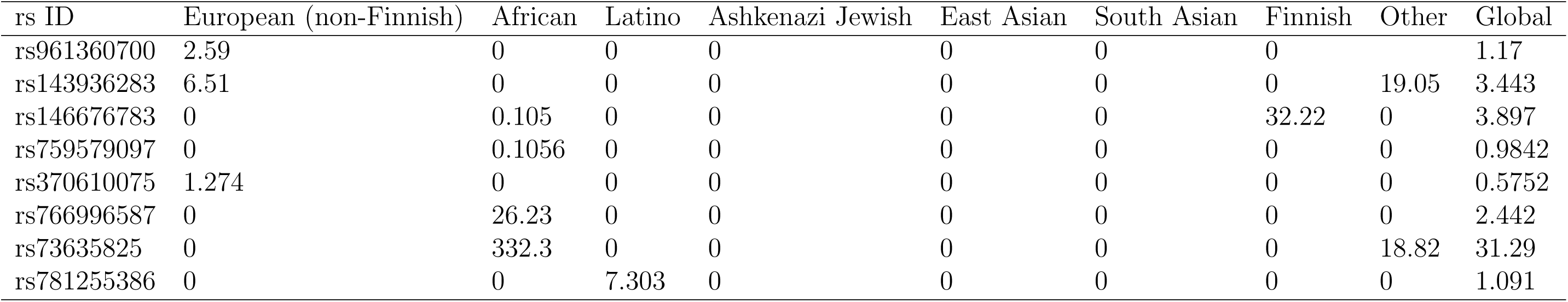
Population frequencies of hACE2 missense variants located on the interaction surface with SARS-CoV-2 RBD (×10−^5^)

| rs ID | European (non-Finnish) | African | Latino | Ashkenazi Jewish | East Asian | South Asian | Finnish | Other | Global |
| --- | --- | --- | --- | --- | --- | --- | --- | --- | --- |
| rs961360700 | 2.59 | 0 | 0 | 0 | 0 | 0 | 0 |  | 1.17 |
| rs143936283 | 6.51 | 0 | 0 | 0 | 0 | 0 | 0 | 19.05 | 3.443 |
| rs146676783 | 0 | 0.105 | 0 | 0 | 0 | 0 | 32.22 | 0 | 3.897 |
| rs759579097 | 0 | 0.1056 | 0 | 0 | 0 | 0 | 0 | 0 | 0.9842 |
| rs370610075 | 1.274 | 0 | 0 | 0 | 0 | 0 | 0 | 0 | 0.5752 |
| rs766996587 | 0 | 26.23 | 0 | 0 | 0 | 0 | 0 | 0 | 2.442 |
| rs73635825 | 0 | 332.3 | 0 | 0 | 0 | 0 | 0 | 18.82 | 31.29 |
| rs781255386 | 0 | 0 | 7.303 | 0 | 0 | 0 | 0 | 0 | 1.091 |

**Figure 4.**
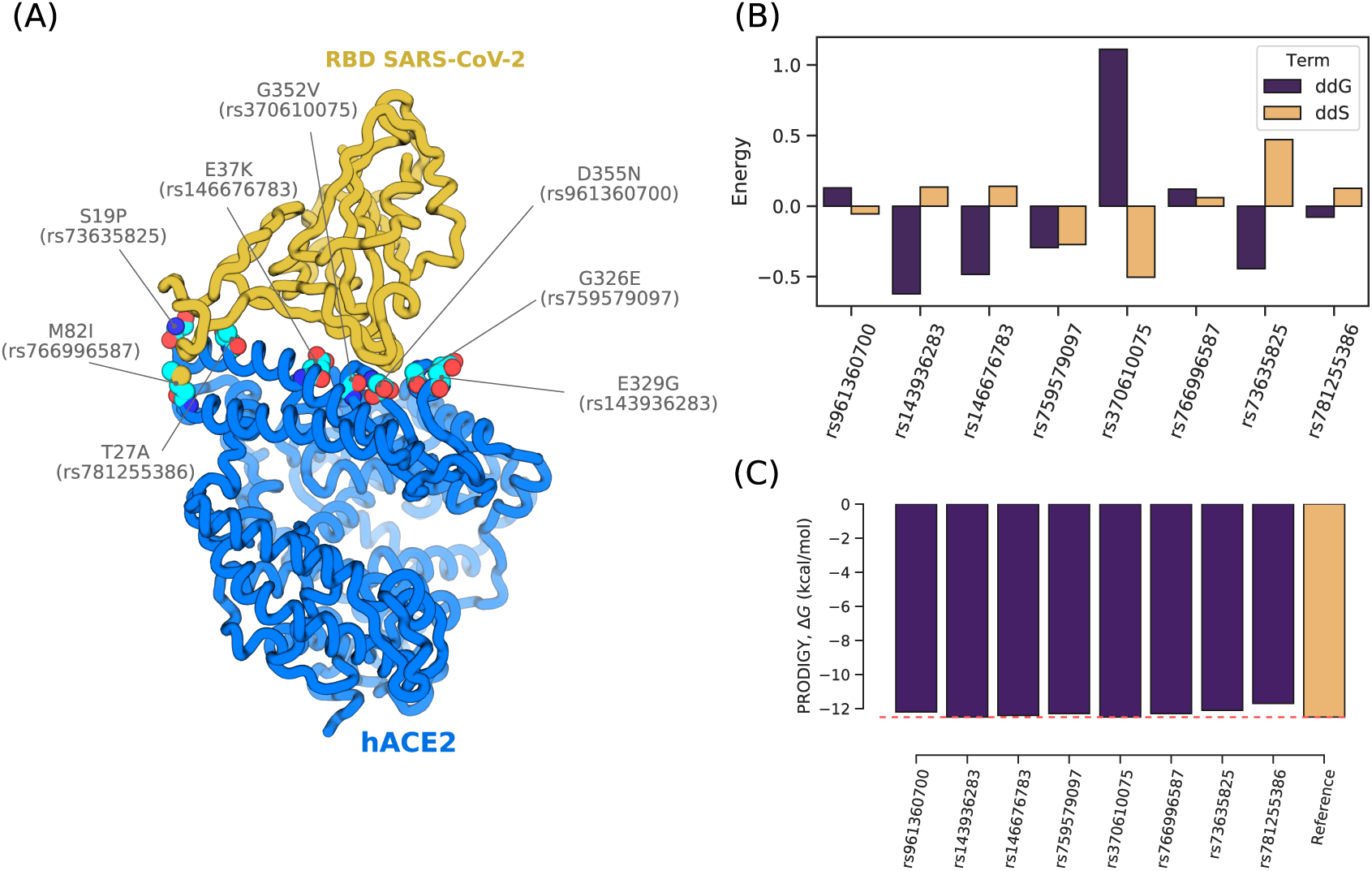
Analyzing the interaction of SARS-CoV-2 RBD with different variants of hACE2. (A) Localization of the variants, labeled by the amino acid change and the dbSNP ID, on the interaction surface of hACE2 and RBD from SARS-CoV-2. Estimation of the changing upon mutation for hACE2 variants calculated for enthalpy (ddG) and entropy (ddS) terms of the folding energy calculated with DynaMut (B) and the interaction energy calculated with PRODIGY (D).

## 4 Discussion

Since the Covid-2019 outbreak, several milestone papers have been published to examine the particularity of SARS-CoV-2 spike protein and its putative interaction with ACE2 as a receptor [21]. In the current study, we focused our analysis on the interface segments of SARS-CoV-2 spike RBD interacting with ACE2 from different species by estimating interaction energy profiles.

We have studied the effect of eight variants of ACE2 in order to detect polymorphisms that may increase or decrease virulence in the host. Our results showed that if ACE2 is the only route for the infection in humans, variants interacting physically with RBD are not likely to disrupt the formation of the complex and would have a marginal effect on the affinity.

Therefore, it is unlikely that any form of resistance to the virus, related to the ACE2 gene, exists. However, this analysis merits to be investigated in depth in different ethnic groups for a better assessment of the contribution of genetic variability in host-pathogen interaction.

The similar values of binding energy scores with different ACE2 orthologues suggest that the ability of binding to different ACE2 orthologous is preserved in many species either for SARS-CoV-2 or SARS-CoV. Therefore, the transition to the zoonotic form is trivial if that depends only on ACE2 as the primary route to the infection in both the intermediate and the final host. However, we know that such a process is very complex since it requires many protein-protein interactions to acquire the specific capacity of infecting and replicating in the host cells [18]. Consequently, it makes sense to assume that many other types of receptors or co-receptors may be critical to determine the capacity of crossing the species barrier. This has been already suggested for SARS-CoV [1] and similarly, SARS-CoV-2 may show the same feature. Moreover, our results show that the significant overlap of glycosylation sites with the protein-protein interface implies a likely interaction of SARS-CoV-2 progenitors with receptors other than ACE2. Finally, recent transcriptomic profiling has suggested the possibility of multiple route infections via the interaction of many human receptors for SARS-CoV-2 [11].

Whole-genome phylogenetic analysis of the different isolates included in this study is consistent with previous works that place the Wuhan-Hu-1 isolate close to Bat-SL-CoVZC45 and Bat-SL-CoVZXC21 isolates [10, 17] within the Betacoronavirus genus. The use of RBD sequences, however places the virus in a clade that comprises SARS-CoV related homologs including isolates from Bat and Civet. The clade swapping as seen in figure 1A, seems also to occur for RaTG13 and Rm1 isolated from bat. This is expected as the use of different phylogenetic markers may considerably affect the topology of the tree. However, The significant divergence in the interfaces segments as a key molecular element contributing to the determination of the tree topology has driven our work toward studying their impact on the interaction with hACE2. The binding of the spike glycoprotein to ACE2 receptor requires a certain level of affinity. In the case where the RBD evolves from an ancestral form closer to that of Bat-SL CoVZC45 and Bat-SL-CoVZXC21, we expected a decrease of the bindin energy scores through the evolution process following incremental changes in the RBD. In such a scenario, we presume that there are other intermediar forms of coronavirus that describe such variation of the binding energ score to reach a level where the pathogen can infect humans with high affinity toward hACE2. On the other hand, our results show that the binding energy score and the interface sequence of SARS-CoV-2 RBD are closer to SARS-CoV related isolates (either from Human or other species). Therefore a recombination event involving the spike protein that might have occurred between SARS-CoV and an ancestral form of the current SARS-CoV-2 virus might be also possible. This will allow for the virus to acquire a minimum set of residues for the interaction with hACE2. The recombination in the spike protein gene has been previously suggested by Wei et al in their phylogenetic analysis [4]. Thereafter, incremental changes in the binding interface segment will occur in order to reach a better affinity toward the receptor. One of these changes may involve P499 residue which substitution to threonine seems to drastically destabilize the interface segment and has a distant effect. Moreover, the decomposition of the interaction energy showed that 5 out of 6 hotspot amino acids in SARS-CoV-2 have their equivalent in SARS-CoV including N501. Contrary to what Wan et al [17] have stated, the single mutation N501T does not seem to enhance the affinity. Rather, the residue Q493 might be responsible for such higher affinity due to a better satisfaction of the Van der Waals by the longer polar side chain of asparagine. Indeed, when we made the same analysis while mutating Q493 to N493, the favorable contribution decreases from −2.55 kcal/mol to a non significant value of −0.01 kcal/mol, thus supporting our claim.

No major divergence of the interaction interface of SARS-CoV-2 RBD with hACE2 was noticed from the similarity matrix analysis. This suggests that the molecular elements required for the binding with the receptor might also be involved in the interaction with other orthologous forms of ACE2 and that these elements are not optimized specifically for the human form. Therefore, it is unlikely that the interface of RBD from SARS-CoV-2 is a result of human intervention via genetic engineering aiming to increase the affinity toward ACE2. For example, residue E484 contributes unfavorably to the binding energy with 2.24 kcal/mol due to an electrostatic repulsion with E75 from hACE2. This residue is an apparent choice for engineering a protein-protein complex with high affinity by substituting E484 with a polar residue. It is, however, noteworthy that the lesser homogeneity of the nodes of SARS-CoV-2 group, in comparison to SARS-CoV, may suggest a higher tolerance for the mutation of the new virus which would allow it to cross the species barrier more easily and to efficiently optimize the interaction in the host.

## Acknowledgement

This work was supported by the South African National Research Foundation (NRF) and the Tunisian Ministry of Higher Education and Scientific Research. The author H. Othman would like to thank Shahine Othman for being understanding about his absence and for not being able to bring him the marshmallow candy because of the COVID-19 outbreak.

