## Supplementary data 2 for "Interaction of the spike protein RBD from SARS-CoV-2 with ACE2: similarity with SARS-CoV, hot-spot analysis and effect of the receptor polymorphism"

### 1. Description of the data

The translated genomic sequence of the spike protein coronavirus 2 isolate Wuhan-Hu-1 corresponds to QHD43416.1 accession in NCBI protein database. The first amino acid of the corresponding RBD starts at position 338. However, we assigned the first index to 337 (a shift of -1). Wan et al (2020, DOI: 10.1128/JVI.00127-20) made this shift for unknown reasons when in their paper and because we compared our results to theirs we decided to use the same indexing schema.

We provide the 3D structures in the supplemental material (file "`RBD_SARS-CoV-2-hACE2.pdb`" and "`RBD_SARS-CoV-hACE2.pdb`") with the amino acid indexes of both the receptor and ligands used in this study.

The file "aligment_MAFFT_spike.fasta" are the sequences of RBD aligned with MAFFT and used to construct the phylogenetic tree.

The file “`ACE2_homologues.txt`” contains the results of the prediction of the putative N-glycosylation sites on ACE2 from the different species reported in this study.

### 2. Error margin of PRODIGY, MM-GBSA, and FoldX

Error margins considedred in the analysis of the scoring energies are dedused from previous benchmarking studies: +/-1.89 kcal/mol for PRODIGY (Xue et al, 2016 PMID 27503228), +/-2.31 kcal/mol for FoldX (Xiong *et al*, 2016 PMID 27899282) and +/-15 kcal/mol for MMGBSA as an approximative error (Aldeghi *et al* 2017, PMID 28786670).

Differences in energy between SARS-COV-2 and Bat-SL-CoVZC45/ Bat-SL-CoVZXC21 interacting with ACE2.

- MM-GBSA energy score difference between SARS-COV-2 and Bat-SL-CoVZC45: 19.6 kcal/mol
- MM-GBSA energy score difference between SARS-COV-2 and Bat-SL-CoVZXC21: 36.4 kcal/mol
- PRODIGY energy score difference between SARS-COV-2 and Bat-SL-CoVZC45: 1.8 kcal/mol
- PRODIGY energy score difference between SARS-COV-2 and Bat-SL-CoVZXC21: 1.5 kcal/mol
- FoldX energy score difference between SARS-COV-2 and Bat-SL-CoVZC45: 10 kcal/mol
- FoldX energy score difference between SARS-COV-2 and Bat-SL-CoVZXC21: 11.99 kcal/mol

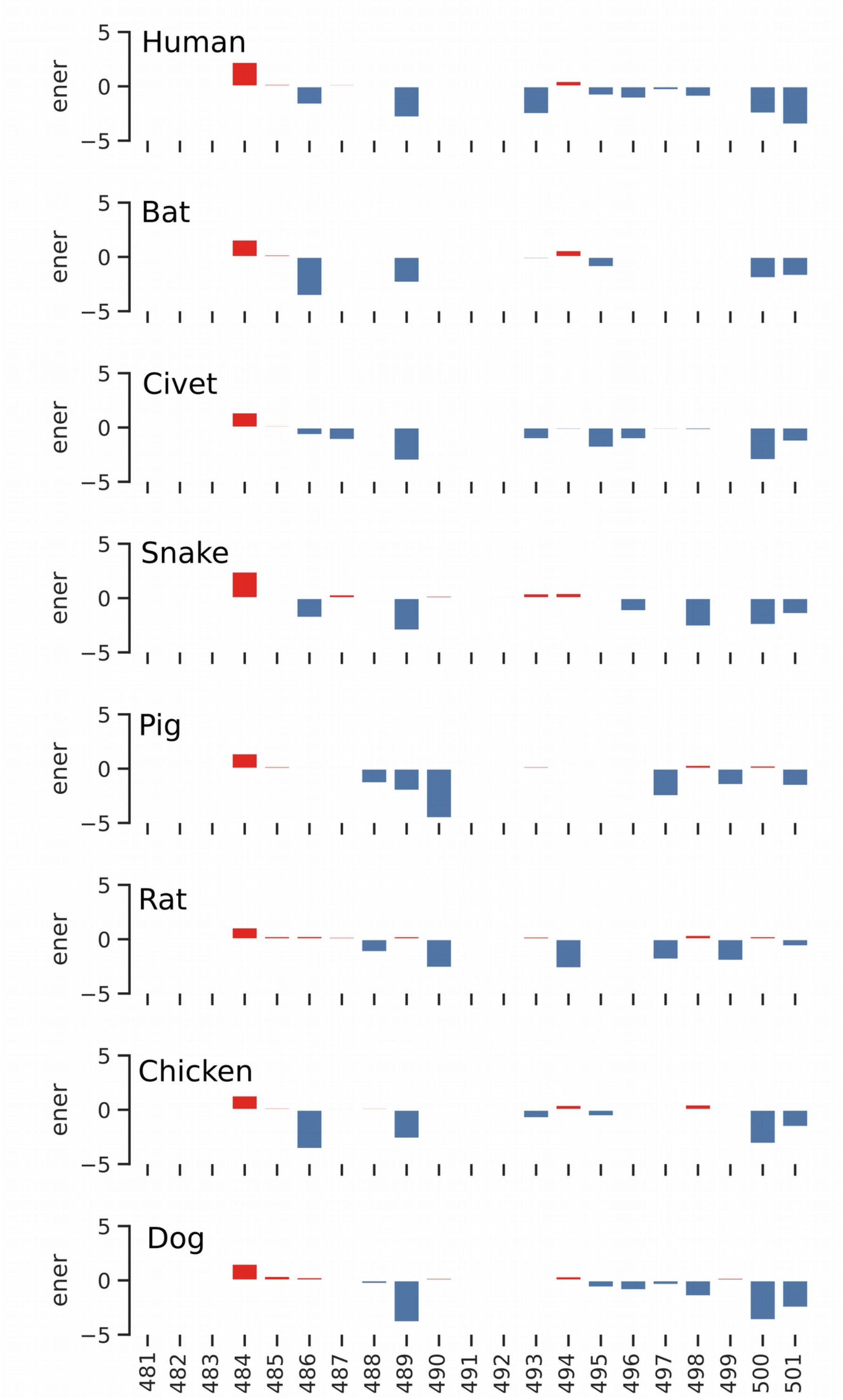


**Figure 1.** Decomposition profiles of RBDs interface residues of SARS-CoV-2 in complex with ACE2 forms from different animals. The profiles were used to calculate a similarity matrix were each specie is assigned to a vector of N components, and N being the number of residues in the interface.

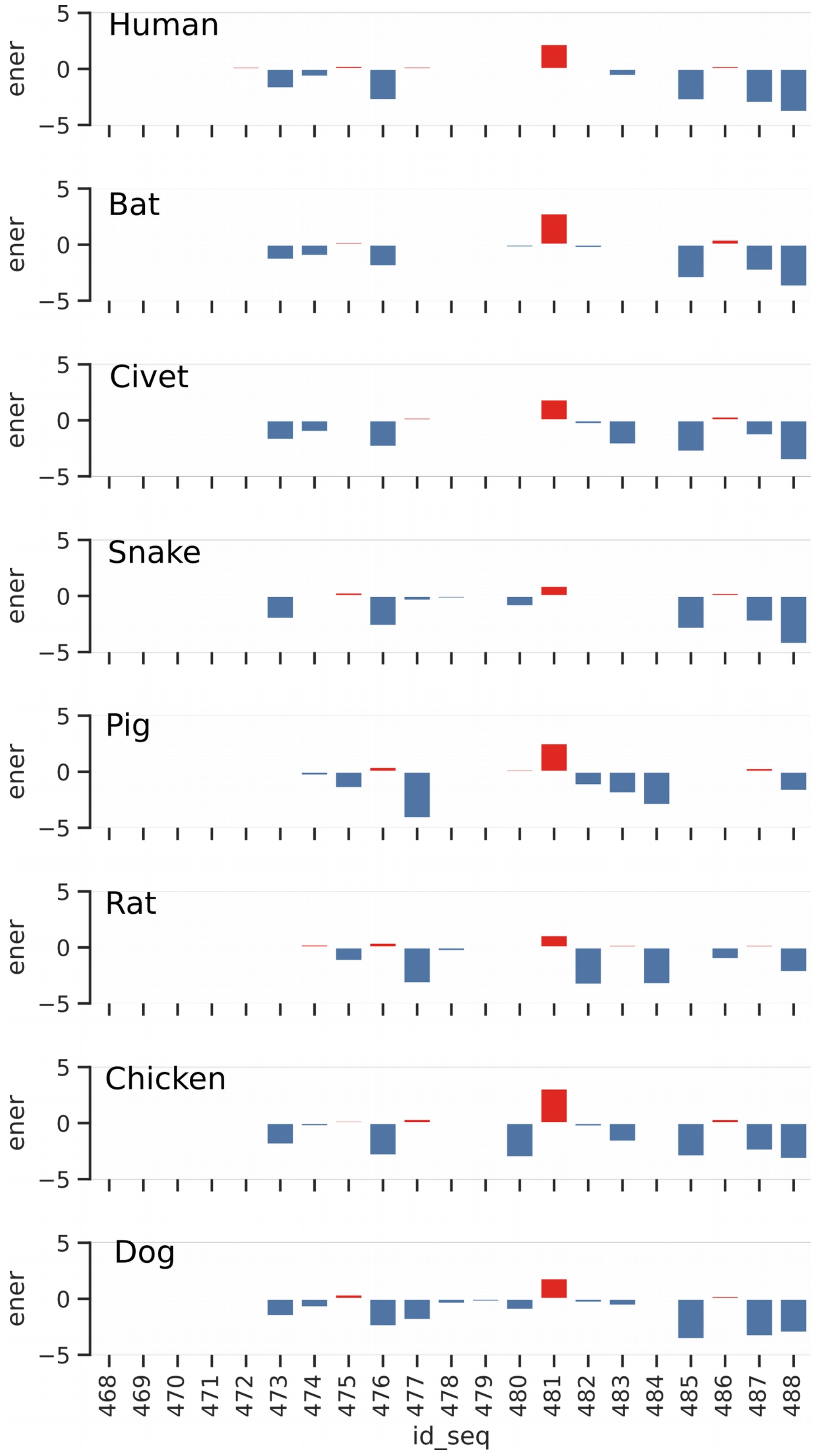


**Figure 2.** Decomposition profiles of RBDs interface residues of SARS-CoV in complex with ACE2 forms from different animals. The profiles were used to calculate a similarity matrix were each specie is assigned to a vector of N components, and N being the number of residues in the interface.

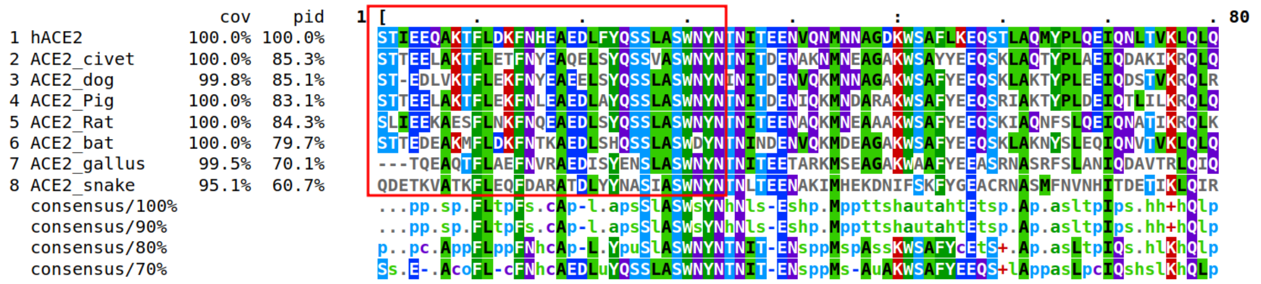


**Figure 3.** Multiple sequence alignment of different ACE2 sequences. The red box indicates the position of the interface segment with RBD.

### 3. Analysis of the putative glycosylation sites

Complexes involving ACE2 proteins from human, civet, dog, chicken, and pig showed no sites of N-glycosylation that overlap with the interface. There is a possibility of steric hindrance caused by residue N82 from bat and rat ACE2 that can alter the binding to the receptor (Figure 4 of supplementary data 2). We noticed however that the snake ACE2 shows a significant overlap of its protein-protein interface with two putative N-glycosylation sites N68 and N105 with the latter predicted with a high degree of confidence.

The rat ACE2 has been shown to have no affinity in binding SARS-CoV (Li et al, 2004, PMID 15452268) due to the presence of Glycan group at position N82. The same outcome would be expected for Rhinolophus sinicus ACE2. This is another indicator that the bat progenitor forms are not optimized for the interaction with human ACE2 since the binding depends on the presence or absence of glycosylated N82. It is also expected that the snake form would not bind to RBD of either SARS-CoV-2 and SARS-CoV.

Next, we have made an examination of the literature as well as some available structures in the PDB. First, in their experiment, Li *et al* (2005, PMID 15791205) have shown that the mutation of N90 to a methionine has enhanced the binding of the pseudotyped lentiviruses with SARS-CoV S protein. This, however, does not influence the geometry of the complex as we will explain further. The same co-Crystal structure of SARS-CoV/hACE2 solved by Li *et al* in 2005 (PMID 16166518) shows a glycosylated N90. The unit cell contains two complexes. In the first, four glycan monomers have been assigned to the density map, none of them, however, was able to establish contacts with RBD as the closest distance to the ligand is of 4.1 Angstroms. In the second complex of the unit cell, only one glycan monomer is assigned to the electron density map and it has a different conformation compared to the glycan group in the other protein-protein complex in the co-Crystal but did not establish any contact with RBD residues. The same observations were noticed for the recently published structure of SARS-CoV-2 RBD with hACE2 in which the glycan groups have different conformations in the two complexes from the unit cell (Shang et al, PMID 32225175). This shows that the N90 glycosyl is highly flexible and therefore would not have a stable contact with RBD from SARS-CoV or SARS-CoV-2. Also, since the twin complexes in the unit cell are very similar (RMSD of 1.2 Angstroms in SARS-CoV2/hACE2 and 1.9 Angstroms in SARS-CoV/hACE2 complexes), the glycosylation, therefore, would not affect significantly the binding mode of RBD in term of geometry (positioning of the atoms of the partners toward each others). Based on all these pieces of evidence, we believe that our results are still reliable even without considering the glycosylation state of this residue.

As for the RBD of SARS-CoV-2 (spanning the segment 337-516), recent efforts by Watanabe *et al* (2020, https://doi.org/10.1101/2020.03.26.010322 ) showed that only residue N343 is glycosylated. This amino acid is situated far from the interface at the first alpha-helix at the N-termini. It is unlikely that glycosylation at this level would introduce a sterical hindrance to the formation of the complex.

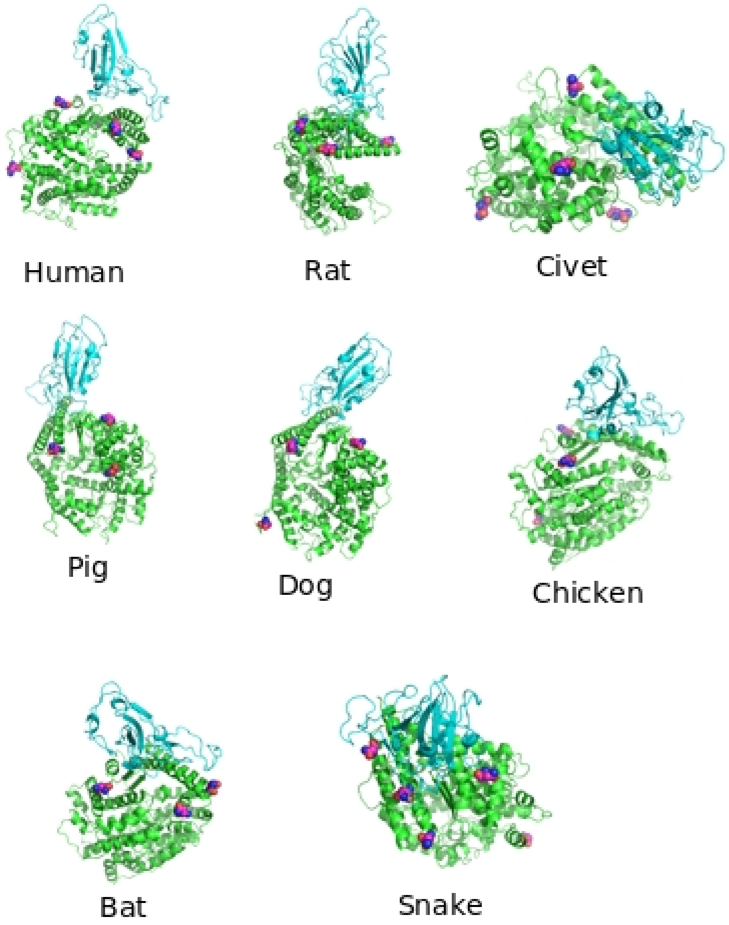


**Figure 4. Location of the putative N-glycosylation sites (magenta) on ACE2 interacting with RBD-SARS-CoV-2.**
